# Learning with Recurrence Geometric AI in Spatial Transcriptomics

**DOI:** 10.64898/2026.08.02.742287

**Authors:** Tuan D. Pham

## Abstract

Cross-domain analysis of spatial transcriptomics is challenging because tissues from different organs, diseases and experimental platforms exhibit distinct cellular compositions, spatial organisations and technical biases, making direct comparison of tissue states difficult. Existing methods primarily focus on domain integration or batch correction but generally do not explicitly model the intrinsic geometry underlying tissue-state organisation across biological systems. This paper presents recurrence geometric artificial intelligence (RGAI), a geometric deep-learning framework for discovering and aligning latent tissue states across heterogeneous spatial transcriptomic domains. RGAI first learns domain-specific latent representations using variational graph autoencoders while simultaneously estimating a Riemannian metric tensor that captures the local geometry of each latent manifold. Geodesic distances induced by the learned metric are used to construct multiscale recurrence graphs that characterise intrinsic tissue-state organisation independently of the original measurement space. Cross-domain manifold correspondence is then established through entropy-regularised Gromov–Wasserstein alignment, after which fuzzy clustering identifies latent tissue states and optimal transport aligns tissue-state signatures across domains. Evaluation on six human spatial transcriptomic datasets spanning wound healing, periodontitis, oral squamous cell carcinoma, head and neck squamous cell carcinoma, cardiac tissue and colorectal cancer shows that RGAI automatically determines biologically meaningful latent tissue-state complexity and identifies coherent recurrence-based tissue states within each domain. The learned geometric representations enable cross-domain alignment of latent manifolds while preserving biologically interpretable tissue-state correspondences despite substantial differences in cellular composition and tissue architecture, demonstrating that integrating learned Riemannian geometry, recurrence analysis and optimal transport provides a robust and interpretable framework for cross-domain tissue-state discovery and comparison in spatial transcriptomics.

## 1 Introduction

Spatial transcriptomic profiling makes it possible to measure gene expression while preserving each measurement’s location within intact tissue, revealing how cellular programmes are organised across the histological structures that molecular dissociation destroys [1, 2, 3]. This capability has been applied across pathologies—skin wound repair [4], periodontal inflammation [5], oral and head-and-neck carcinoma [6], cardiac injury [7], and colorectal tumours [8]—almost always as an independent study with its own patients, platform, and processing pipeline. A natural question follows: do seemingly distinct pathologies— an inflamed gingival margin, a healing dermal wound, a fibrotic myocardium, a tumour stroma—share conserved underlying tissue-state programmes (fibrosis, chronic inflammation, angiogenesis, epithelial-mesenchymal transition), or are the regularities within each dataset artefacts of that dataset alone? Answering this requires a framework that can discover disease-relevant tissue states *within* each dataset and then establish principled correspondences *across* datasets that were never designed to be compared.

That combination is harder than either task alone. Within a single dataset, tissue states rarely form crisp, well-separated clusters: inflammation grades continuously into resolution, granulation tissue grades into scar, and a spot at the boundary between two programmes is genuinely mixed rather than misclassified. Across datasets, there is no natural correspondence to exploit—different patients, platforms, and numbers of profiled locations—so comparison must proceed without paired examples, matching structure to structure rather than observation to observation. An adequate framework therefore needs a notion of tissue-state similarity that respects the nonlinear geometry of expression space, a mechanism for graded rather than hard state membership, and a principled way to align discovered structure across domains that were never jointly measured.

Each of these requirements points toward a different, independently mature body of work. Representing a learned latent space as a Riemannian rather than a Euclidean manifold, so that distances adapt to local geometry rather than a fixed Euclidean metric, is well established in general representation learning [9, 10, 11, 12, 13] and has already been applied to disease modelling [14]. Treating a biological process as *recurrent* rather than as a single trajectory has a long history in nonlinear dynamics [15], and this paper builds directly on the author’s own recent application of that idea to cross-domain tissue-state conservation [16, 17]. Expressing state membership as continuous and graded is the founding idea of fuzzy clustering [18]. Discrete curvature—how locally convergent or divergent a neighbourhood is under a given metric—is a useful signature of stability versus transition in biological and other complex networks [19, 20, 21]. Aligning structure across domains without paired correspondences is precisely the problem optimal transport was built for [22, 23, 24, 25], and Gromov–Wasserstein (GW) transport specifically has already been used to align single-cell datasets that share no common coordinate system [26].

Artificial intelligence—specifically, end-to-end trainable neural representation and metric learning—lets these four ingredients operate jointly rather than as separately fitted stages. Domain-specific encoders, a shared geometric encoder, and a metric network jointly learn the shared and domain-specific latent coordinates and the local Riemannian metric, with additional supervision from a biological-annotation head where labels are available; the recurrence tensor and curvature descriptors are then derived from this learned geometry rather than fixed in advance or hand-designed. This is the sense in which the framework constitutes an AI system rather than a classical geometric construction.

What has not been done is to combine these four ingredients into a single, jointly trained framework aimed at the question this paper opened with: whether tissue-state programmes discovered independently within multiple, disparately sourced spatial transcriptomic datasets correspond to one another in a geometrically principled sense. This paper introduces Recurrence Geometric AI (RGAI), a framework that learns a shared, domain-aware latent manifold for spatial transcriptomic data, equips it with a directly parameterised Riemannian metric, and uses the resulting geodesic structure to drive three downstream analyses: multiscale recurrence characterisation of local tissue stability, Ollivier– Ricci curvature as a descriptor of transition versus cohesion, and fuzzy discovery of disease-relevant tissue states. States discovered independently within each domain are then aligned across domains through a two-stage optimal-transport procedure—coarse, landmark-based GW alignment of the manifolds themselves, followed by entropy-regularised transport between the discovered states, using a composite cost combining geodesic distance, curvature agreement, transition-behaviour similarity, and biological-annotation similarity where available.

This paper makes the following contributions:

- A jointly trained encoder–metric architecture (Section 3.4) learning domain-shared and domain-specific latent representations together with a directly parameterised Riemannian metric tensor, so distances adapt to the local geometry of the latent manifold.
- A multiscale, geodesic-based recurrence formalism (Section 3.5) and local differential-geometric descriptors, including Ollivier–Ricci curvature (Section 3.6), characterising local stability, transition, and structural distortion of the learned manifold.
- A recurrence- and geometry-regularised fuzzy clustering scheme (Section 3.8) for disease-state discovery admitting graded, partial membership rather than a hard partition, with an analysis of state-to-state spatial transition behaviour (Section 3.9).
- A two-stage cross-domain alignment procedure (Section 3.10) combining coarse GW manifold alignment with a state-level, entropy-regularised OT correspondence, using a composite cost integrating geometric and biological evidence.
- An empirical evaluation (Section 4) across six independently sourced spatial transcriptomic datasets spanning tissue repair, chronic inflammatory disease, cardiovascular pathology, and epithelial cancers.

The remainder of this paper is organised as follows. Section 2 reviews prior work along the four axes above. Section 3 describes the six datasets and the proposed approach, from representation learning through cross-domain alignment. Section 4 reports empirical results. Section 5 discusses findings, limitations, and future work. Section 6 concludes the study.

## 2 Related Work

RGAI draws on several established lines of work, each addressing one facet of the proposed framework in isolation. This section situates RGAI relative to prior work along four axes: learned Riemannian metrics for biological latent spaces, discrete curvature as a descriptor of cell or tissue state, GW and entropy-regularised transport for cross-domain single-cell alignment, and the previously proposed cross-domain tissue-state framework.

### Learned Riemannian metrics on biological manifolds

Treating a learned latent space as Riemannian rather than Euclidean is well established in representation learning [9, 10, 11] and has been applied to biological data: Gruffaz et al. [14] learn a Riemannian metric, expressed as the push-forward of the Euclidean metric under an estimated diffeomorphism, to model patient-specific disease-progression trajectories from longitudinal biomarkers. RGAI adopts the same principle but differs in scope: the metric tensor in Section 3.4 is a directly parameterised, per-point Cholesky-style construction rather than a single global diffeomorphism, is learned jointly with a multi-domain, multi-modal encoder rather than fit to one cohort, and feeds downstream curvature and GW alignment terms absent from [14].

### Discrete curvature as a state or trajectory descriptor

Discrete Ricci curvature computed on cell- or gene-similarity graphs has previously characterised biological state: Baptista et al. [21] use Forman–Ricci curvature and a discrete Ricci flow on protein-interaction networks weighted by single-cell transcriptomes to distinguish stem, differentiated, and cancerous states, extending earlier work using Ollivier–Ricci curvature on cancer networks [19, 20]. RGAI’s use of Ollivier–Ricci curvature (Section 3.6) targets the same idea— curvature as a proxy for local stability versus transition—but computes it on a *k*-nearest-neighbour graph of the *learned* latent manifold under the *learned* metric of Section 3.4, rather than a fixed network with hand-specified weights, treating curvature as one of several descriptors feeding a downstream fuzzy state model rather than a flow interpolating between two fixed endpoints.

### GW alignment across domains

Cross-domain alignment via GW optimal transport is itself established in single-cell genomics: SCOT [26] aligns single-cell multi-omics datasets by matching the intra-domain graph-distance structure of each dataset through entropic GW transport, without paired correspondences. The coarse alignment loss in Section 3.10 uses the same GW formalism [24] for the same reason—unpaired, structure-preserving matching—but applies it as a periodically-refreshed regularisation target on the encoder and metric-network parameters throughout training, evaluated on resampled landmark subsets, rather than a one-off post hoc alignment of fixed embeddings. RGAI additionally performs a second, independent entropic OT matching at the level of discovered fuzzy disease states (also Section 3.10), using a composite cost combining geodesic distance, curvature, transition-distribution divergence, and biological-annotation similarity; this state-level step has no counterpart in SCOT, which aligns individual cells rather than discovered states.

### Relationship to the previous cross-domain tissue-state framework

RGAI extends the previously proposed cross-domain tissue-state framework [16], which integrates spatial transcriptomic data across wound healing, oral, and cardiovascular tissues using recurrence modelling, graph-based spatial learning, fuzzy tissue-state analysis, and tensor decomposition to identify conserved fuzzy tissue states. RGAI builds directly on this earlier recurrence-based, fuzzy-state, cross-domain perspective, and its principal methodological advances are the differential-geometric formalisation of the latent representation, including an explicitly learned Riemannian metric and its geodesics (Section 3.4); Ollivier–Ricci curvature as an explicit geometric descriptor (Section 3.6); and the replacement of heuristic cross-domain matching with GW manifold alignment plus a separate entropy-regularised optimal-transport procedure for tissue-state correspondence (Section 3.10).

## 3 Methods

### 3.1 Datasets

RGAI was evaluated using six publicly available human spatial-transcriptomic datasets spanning tissue repair, chronic inflammation, cancer, and cardiovascular disease, chosen to test whether the geometric representation could identify shared tissue-state organisation across biologically distinct tissues. All datasets were obtained from GEO and comprised 10x Genomics Visium data: human cutaneous wound healing (GSE241124), periodontitis (GSE206621), oral squamous cell carcinoma (OSCC; GSE208253), head and neck squamous cell carcinoma (HNSCC; GSE181300), cardiac tissue from healthy and failing hearts (GSE135805), and primary colorectal cancer (CRC; GSE226997).

The wound-healing dataset captures the inflammatory, proliferative, and remodelling phases of human skin repair [4]. The periodontitis dataset profiles healthy and chronically inflamed gingival tissues, capturing spatially localised inflammatory fibroblast and immune-cell programmes [5]. The OSCC dataset comprises tumour-core and invasive-margin sections from HPV-negative oral tumours [6], and the HNSCC dataset contains primary tumour sections from spatially distinct intratumoural regions [27]. The cardiac dataset includes donor hearts together with explanted hearts affected by hypertrophic, dilated, or ischaemic cardiomyopathy [7], and the CRC dataset contains Visium profiles from primary colorectal tumours capturing spatial variation in malignant, stromal, and immune compartments [8]. Together these datasets span substantial diversity in tissue architecture, disease mechanisms, spatial organisation, sample size, spot density, and transcriptional composition (Table 1).

**Table 1.** Spatial transcriptomic datasets used to evaluate RGAI.

| **Dataset** | **GEO accession** | **Disease/tissue** | **Platform** |
| --- | --- | --- | --- |
| Wound healing | GSE241124 | Human cutaneous wound healing | 10x Visium |
| Periodontitis | GSE206621 | Chronic periodontitis | 10x Visium |
| OSCC | GSE208253 | Oral squamous cell carcinoma | 10x Visium |
| HNSCC | GSE181300 | Head and neck squamous cell carcinoma | 10x Visium |
| Cardiac | GSE135805 | Healthy and failing human hearts | 10x Visium |
| CRC | GSE226997 | Primary colorectal cancer | 10x Visium |

For each specimen, the spot-level gene-expression matrix and spatial coordinates were retained. Standard quality control removed low-quality observations and rarely detected genes before library-size normalisation and log transformation, and highly variable genes were identified independently within each dataset to preserve domain-specific variation prior to alignment. No histopathological annotations, tissue labels, or clinical outcome information were used during training or state discovery; where available, such annotations were reserved for post hoc interpretation and validation.

### 3.2 Proposed Approach

RGAI integrates four complementary components: (i) AI for learning latent biological representations from heterogeneous biomedical data [16, 17]; (ii) differential geometry [12, 13] for endowing the latent manifold with a learned Riemannian metric defining intrinsic distances, curvature, and local structure; (iii) recurrence analysis [15, 16] for identifying stable, recurrent, and transitional disease states on the manifold; and (iv) optimal transport [22, 23] for aligning these geometric state representations across domains, together providing a mathematically principled framework for discovering conserved disease-state organisation beyond individual diseases and measurement technologies.

The central premise is that heterogeneous biomedical observations lie on a lower-dimensional latent manifold whose intrinsic geometry reflects biologically meaningful state organisation. RGAI jointly learns latent representations, local geometric structure, fuzzy recurrence relationships, disease-state memberships, and cross-domain correspondences.

Let 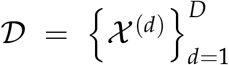 denote a collection of *D* biological domains, where 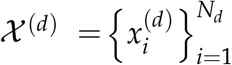 contains *N*_*d*_ observations from domain *d*, each the gene-expression profile measured at a single spatial transcriptomic location. Each domain is associated with a structural graph 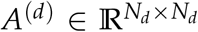, encoding spatial adjacency, temporal ordering, specimen structure, or nearest-neighbour relationships. *N* (*i*) denotes the neighbours of observation *i* under *A*^(*d*)^ (or, where noted, under a *k*-nearest-neighbour graph in latent space), and *E*^(*d*)^ the corresponding edge set.

### 3.3 Shared and Domain-Specific Representation Learning

Each observation is first mapped to an intermediate domain-specific representation,

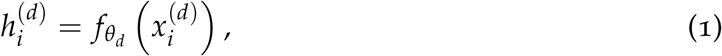

where 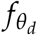 is a domain-specific encoder. A shared geometric encoder then projects the intermediate representation onto a common latent manifold ℳ,

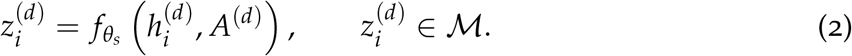

To distinguish conserved biological structure from domain-specific variation, the latent representation is decomposed as

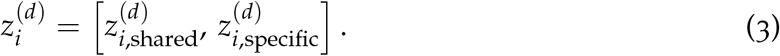

The shared component captures recurrent state organisation across diseases, whereas the domain-specific component retains variation unique to each dataset. This decomposition is not automatic: without an explicit mechanism, nothing prevents domain-specific information from leaking into 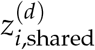 , or vice versa. A disentanglement penalty in the form of a soft subspace-orthogonality constraint is therefore introduced [28],

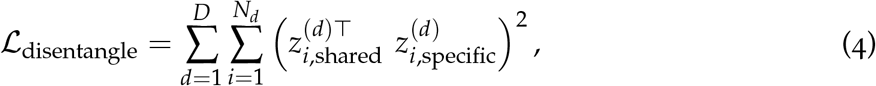

which penalises linear dependence between the shared and specific subspaces at every observation, encouraging the two embeddings toward orthogonality and promoting a decomposition in which the shared representation captures domain-invariant characteristics while the domain-specific representation encodes features unique to each domain.

### 3.4 Learning the Local Riemannian Metric

Rather than assuming that the latent manifold is Euclidean, RGAI learns a local positive-definite metric tensor [9, 10, 11],

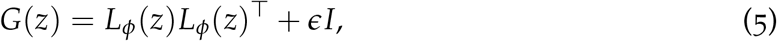

where *L*_*ϕ*_ is a neural network parameterised by *ϕ* mapping each latent point 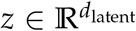 to a matrix 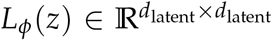, learned jointly with the remaining components of RGAI so that the metric adapts to the local organisation of the latent representations. The entries of *L*_*ϕ*_(*z*) determine the local scaling of individual latent directions and their interactions. Although *L*_*ϕ*_(*z*) need not itself be symmetric or positive definite, the product *L*_*ϕ*_(*z*)*L*_*ϕ*_(*z*)^⊤^ is necessarily symmetric and positive semidefinite. Here 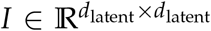 is the identity matrix and *ϵ >* 0 is a small constant ensuring *G*(*z*) is strictly positive definite and numerically stable.

For neighbouring latent points *z*_*i*_ and *z*_*j*_ with (*i, j*) ∈ *E*^(*d*)^, the local geometric distance is approximated by

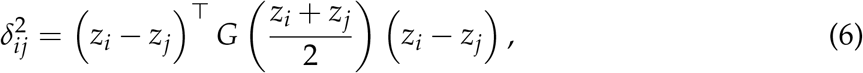

and the corresponding graph edge weight is 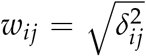. Because *δ*_*ij*_ is computed directly from *z*_*i*_, *z*_*j*_, and *G*, it is smooth and fully differentiable in the encoder and metric-network parameters [9].

For pairs of latent points not directly connected in the graph, the intrinsic geodesic distance is approximated, as in Isomap [29], by the shortest path through the weighted graph *W*^(*d*)^ = (*w*_*ij*_),

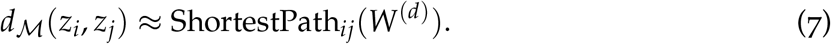

#### Remark 1

(Differentiability of geodesic distances). *Unlike δ*_*ij*_, *the shortest-path operator in Eq*. (7) *is a discrete combinatorial function of W*^(*d*)^ *and is not differentiable with respect to the encoder or metric-network parameters in the ordinary sense*.

To keep the learned metric well behaved, a metric regularisation loss combines two terms: an anti-degeneracy term discouraging *G* from collapsing towards singularity or excessive distortion, and a smoothness term discouraging abrupt variation between neighbouring latent points:

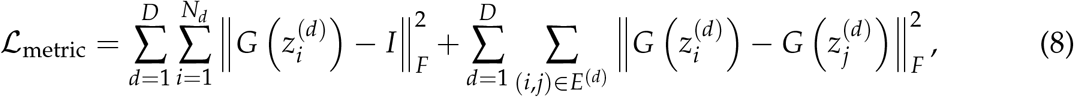

where ∥ · ∥_*F*_ denotes the Frobenius norm and *I* is the identity matrix. The first term biases the metric towards Euclidean unless supported by the data, preventing ill-conditioned or nearly singular tensors; the second enforces local geometric consistency by encouraging neighbouring latent points to have similar metric tensors, reducing spurious fluctuations caused by noise. The learned metric can thus emphasise biologically informative directions while suppressing weakly informative or technical variation, without sacrificing numerical stability or differentiability.

### 3.5 Multiscale Geometric Recurrence

The recurrence relationship between two observations is defined in terms of their geodesic distance on the learned manifold. At scale *s*, the geometric recurrence strength is

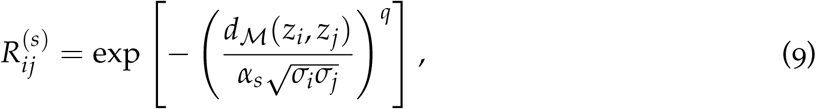

where *α*_*s*_ *>* 0 determines the recurrence scale, with smaller values emphasising local recurrence and larger values capturing broader manifold structure; *q >* 0 controls the decay profile of the recurrence kernel, and *σ*_*i*_ *>* 0 is a self-tuning local bandwidth [30], defined as

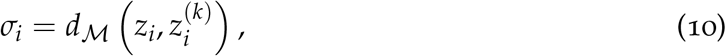

where 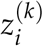 denotes the *k*th nearest neighbour of *z*_*i*_ under geodesic distance. In implementation, *σ*_*i*_ is lower-bounded by a small constant *ε* to avoid numerical instability when observations nearly coincide.

For an ordered set of recurrence scales 0 *< α*_1_ *<* · · · *< α*_*S*_, the recurrence matrices are stacked to form the multiscale recurrence tensor

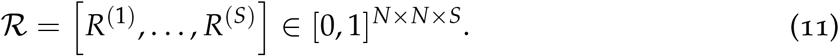

Because the geodesic distance is symmetric and non-negative, the recurrence matrices satisfy

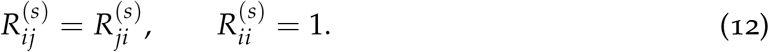

Moreover, for fixed local bandwidths,

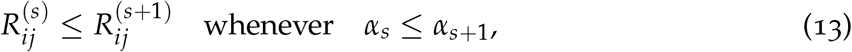

so that progressively larger scales capture increasingly broad regions of the learned manifold.

For spatial transcriptomic data, geometric recurrence is modulated by a spatial kernel,

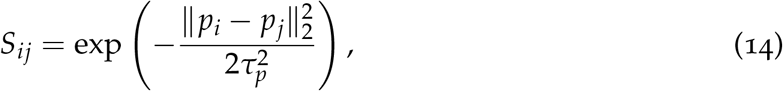

where *p*_*i*_ and *p*_*j*_ denote the spatial coordinates of observations *i* and *j*, respectively, and *τ*_*p*_ *>* 0 is the spatial bandwidth. The resulting spatially aware recurrence strength is

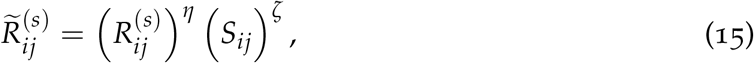

where *η, ζ >* 0 control the relative influence and sharpness of the geometric and spatial components. Equivalently,

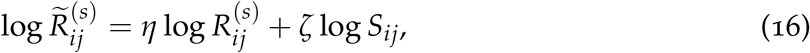

so *η* and *ζ* act as log-linear weights, optionally normalised so that *η* + *ζ* = 1. Because 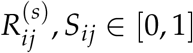, the combined recurrence satisfies 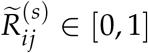 for any *η, ζ >* 0. The spatially aware recurrence matrices are stacked analogously to form

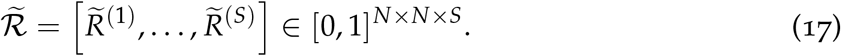

The recurrence tensors are computed from the learned manifold at periodic evaluation checkpoints during training (Remark 1) and, because *d*_ℳ_ involves a discrete shortest-path operation, are not directly differentiated through during optimisation. Geometric supervision at each gradient step is instead provided by the differentiable local-distance, neighbourhood-preservation and metric-regularisation terms.

The multiscale organisation of the recurrence tensor is characterised by the following consistency criterion:

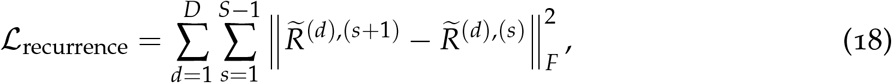

where 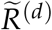 denotes the spatially (or temporally) modulated recurrence tensor (Eq. (15)). To avoid repeatedly recomputing multiscale geodesic recurrence matrices during optimisation, 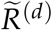 is updated only during periodic geometry-refresh steps, and the consistency criterion is re-evaluated after each refresh using the updated representation. This formulation requires no external supervision or known cross-observation correspondences and provides a quantitative measure of multiscale smoothness in the learned recurrence representation.

### 3.6 Local Differential-Geometric Descriptors

RGAI characterises the local geometry of each observation using complementary measures of recurrence, distortion, anisotropy, and curvature.

The local recurrence density of observation *i* in domain *d* is defined as

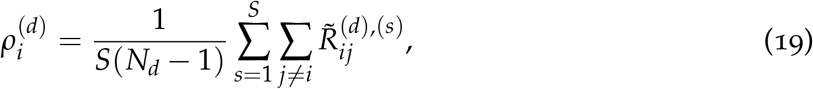

where *S* is the number of recurrence scales and *N*_*d*_ is the number of observations in domain *d*. Averaging over scales provides a scale-aggregated summary of the local recurrence structure while retaining information from the learned multiscale representation.

The local Riemannian volume distortion is measured by the volume element induced by the metric tensor [31]:

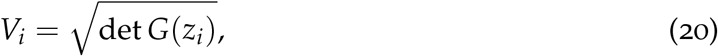

which quantifies local expansion or contraction of the latent space under the learned metric.

Let *λ*_*i*,1_, … , *λ*_*i,m*_ denote the eigenvalues of *G*(*z*_*i*_). Local anisotropy is quantified by the regularised spectral condition number of the metric tensor [35],

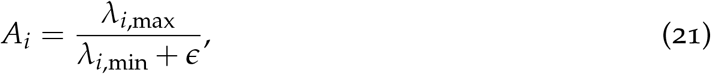

where *λ*_*i*,max_ and *λ*_*i*,min_ are the largest and smallest eigenvalues of *G*(*z*_*i*_), and *ϵ >* 0 ensures numerical stability. Values near unity indicate isotropic geometry; larger values indicate stronger directional anisotropy.

Curvature is estimated using graph-based Ollivier–Ricci curvature [19, 20]. For each observation *i*, define a lazy random-walk neighbourhood measure with laziness *γ* ∈ [0, 1],

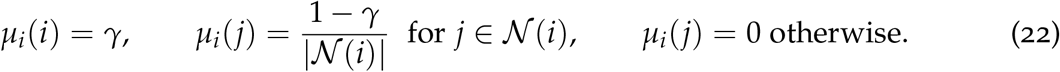

The edge-level curvature between adjacent observations *i, j* is

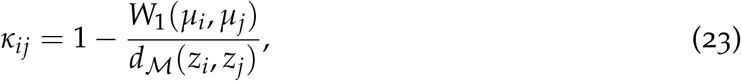

where *W*_1_ denotes the Wasserstein-1 distance [22] between *µ*_*i*_ and *µ*_*j*_ under ground metric *d*_ℳ_. Positive curvature indicates locally cohesive or convergent organisation, whereas negative curvature may indicate branching, divergence, or transition boundaries. Since *κ*_*ij*_ is defined per edge while the descriptor below needs one value per observation, aggregation averages over incident edges,

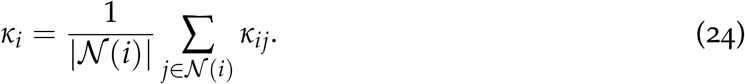

Curvature is not merely a descriptive output but is also used during training to encourage stable manifold geometry, via two regularisation terms: a global term penalizing deviation from the mean curvature 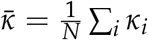, where 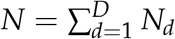 is pooled across *all* domains (curvature is compared globally, unlike the domain-local *N*_*d*_ elsewhere, e.g. Eq. (19), because it lives on the shared manifold ℳ that cross-domain alignment, Section 3.10, seeks to make comparable), discouraging isolated pockets of extreme curvature; and a local term penalising abrupt curvature changes between neighbours, discouraging spurious fragmentation of continuous state regions,

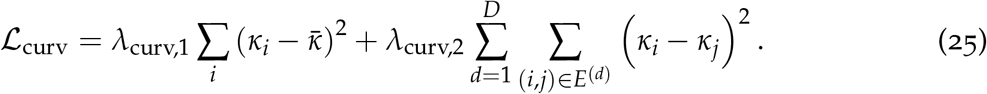

Because *κ*_*ij*_ requires a Wasserstein-1 solve per edge in addition to a geodesic distance, evaluating ℒ_curv_ at every step is at least as costly as the geodesic refresh; it is therefore computed on the same periodic-refresh, stop-gradient schedule as 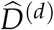 precomputed curvature as fixed targets.

Each observation is therefore assigned a local geometric descriptor

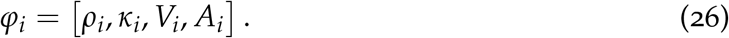

### 3.7 Automatic determination of tissue-state complexity

The number of latent tissue states was determined separately for each domain using a multi-criterion model-selection procedure evaluated across repeated independently trained RGAI models. For each candidate *K*, fuzzy state discovery was applied to every trained recurrence manifold, and the resulting partitions were assessed using cross-run stability, fuzzy-clustering validity and specimen-independence criteria.

Cross-run stability was quantified by the mean pairwise adjusted Rand index (ARI) [32] between hard state assignments from the repeated runs. Fuzzy partition quality was evaluated using the adjusted partition coefficient

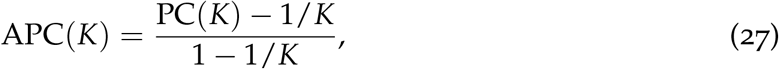

where

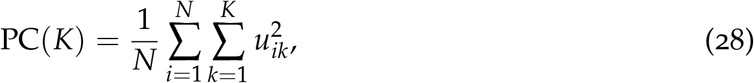

and *u*_*ik*_ denotes the membership of observation *i* in state *k*. Membership uncertainty was measured by the normalised entropy

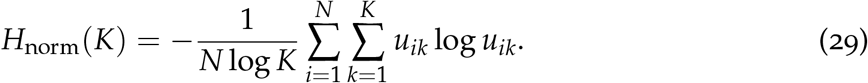

Cluster compactness and separation were assessed using a geodesic Xie–Beni index [33],

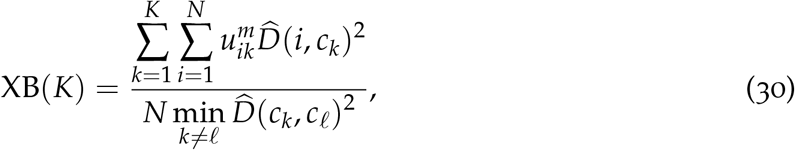

where *c*_*k*_ is the Fréchet (intrinsic) centre [34] of state *k*—the point minimising the weighted sum of squared geodesic distances to observations assigned to that state. To discourage partitions that merely reproduced specimen identity, the normalised mutual information between specimen labels and hard state assignments was also calculated.

For each criterion, candidate values of *K* were ranked across the search range. The aggregate selection score was defined as

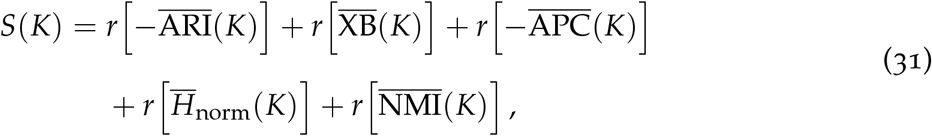

where *r*[·] denotes ascending rank and the overbar denotes the mean across independently trained models. The procedure thus favours high cross-run stability, compact and well-separated states, concentrated fuzzy memberships and weak dependence on specimen identity. Candidate solutions in which one or more states received no maximum-membership assignments were excluded. The selected tissue-state complexity was

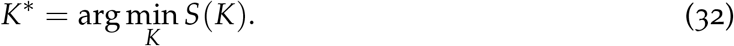

After selection of *K*^∗^, the repeated-run partition with the highest mean ARI to all other partitions was used as the medoid. Membership columns from the remaining runs were matched to the medoid partition and averaged to construct a label-aligned fuzzy consensus, from which state centres, transition matrices, signatures and cross-domain correspondences were recomputed.

### 3.8 Fuzzy Disease-State Discovery

Disease states are represented as fuzzy regions of the learned latent manifold rather than discrete Euclidean clusters. Let *u*_*ik*_ ∈ [0, 1] denote the membership of observation *i* in state *k*, and let **u**_*i*_ = (*u*_*i*1_, … , *u*_*iK*_)^⊤^ be the corresponding membership vector, satisfying the partition constraint

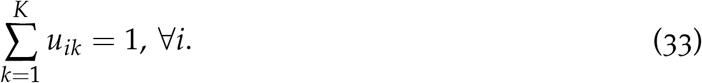

The memberships are estimated by minimising a recurrence- and geometry-regularised extension of the fuzzy *c*-means (FCM) objective [18],

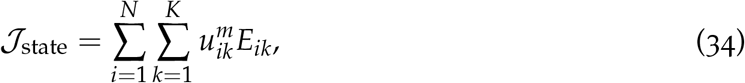

where *m >* 1 is the fuzziness exponent and *E*_*ik*_ denotes the assignment energy of observation *i* to state *k*,

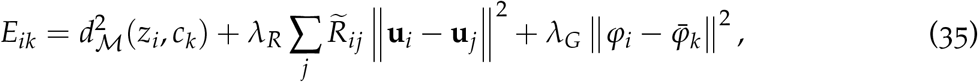

where *c*_*k*_ denotes the Fréchet centre of state *k*, 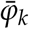 is the mean geometric descriptor of that state, 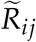 is the multiscale recurrence affinity, and *d*_ℒ_ is the learned geodesic distance. The first term promotes compactness on the manifold, the second encourages recurrence-connected observations to share similar membership vectors, and the third favours geometric consistency within each state, so smaller *E*_*ik*_ corresponds to stronger assignment. When *λ*_*R*_ = *λ*_*G*_ = 0 and *d*_*M*_ is replaced by Euclidean distance, Eq. (34) reduces to the classical FCM objective.

Because the recurrence regularisation couples neighbouring observations through their membership vectors, Eq. (34) is optimised iteratively rather than by a closed-form update, with assignment energies at iteration *t* evaluated from the previous iteration’s memberships and updated according to

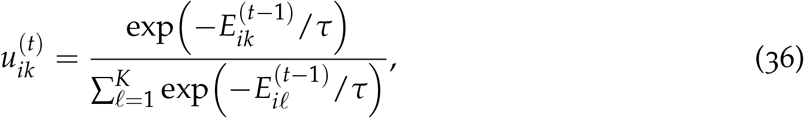

where *τ >* 0 is an annealing temperature. This block-coordinate mean-field update is analogous to deterministic-annealing expectation-maximisation and provides a stable approximation to the coupled optimisation problem.

The centre of each state is computed as a weighted Fréchet mean on the learned manifold. Since the manifold is represented by discrete latent observations connected through the learned geodesic graph, the centre is

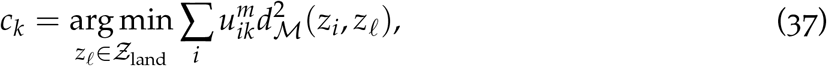

where *Ƶ*_land_ ⊆ *Ƶ* denotes the set of candidate landmark points. This discrete optimization uses the precomputed geodesic distance matrix and is the primary computation throughout RGAI. When higher spatial precision is required, the landmark solution may optionally be refined locally using the learned Riemannian metric,

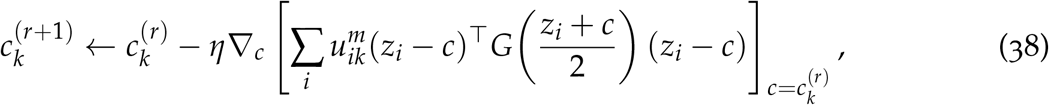

valid within a neighbourhood of the discrete Fréchet centre; this refinement is optional and does not alter the primary optimisation procedure.

### 3.9 Disease-State Transition Analysis

Because all datasets analysed in this study are cross-sectional, relationships between latent tissue states are characterised by spatial adjacency rather than temporal progression. The conditional spatial adjacency probability between states is defined as

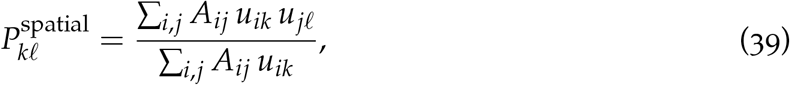

where *A*_*ij*_ is the spatial adjacency matrix, with *A*_*ij*_ = 1 if observations *i* and *j* are neighbouring locations and 0 otherwise. 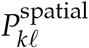 estimates the conditional probability that an observation with high membership in state *k* is spatially adjacent to one with high membership in state *ℓ*, so the resulting matrix summarises the spatial organisation and co-localisation of latent tissue states within each specimen.

Because Eq. (39) is derived solely from spatial neighbourhood relationships, it should not be interpreted as temporal progression or causal state transition. Biological directionality requires additional temporal or mechanistic information—longitudinal sampling, RNA velocity, pseudotime analysis, or independent biological evidence—so here the spatial adjacency matrix is used exclusively to characterise spatial relationships among latent tissue states.

### 3.10 Cross-Domain State Alignment

RGAI aligns geometry across domains at two complementary scales, which are distinguished explicitly below.

#### Coarse, manifold-level alignment (periodic refresh)

A GW discrepancy [24] between pairs of domain manifolds regularises the overall relational geometry towards crossdomain consistency,

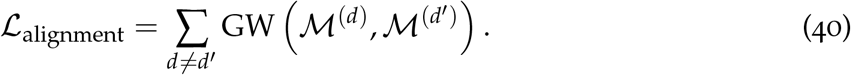

Because GW is computed from pairwise geodesic distances within each domain, it inherits the same non-differentiability as *d*_ℳ_ (Remark 1) and is evaluated on the same periodic-refresh, stop-gradient schedule as the precomputed geodesic distances and curvature, rather than backpropagated through at every step: at each refresh, GW(ℳ ^(*d*)^, ℳ ^(*d*^*′* )) is evaluated on a resampled or fixed landmark subset of *L* ≪ *N*_*d*_ points per domain (exact GW scales quadratically, and its entropic-Sinkhorn relaxation [25] still scales at least quadratically, in points per domain, following standard mini-batch/landmark practice for large-scale optimal transport), and the resulting coupling and discrepancy are held fixed as regularisation targets until the next refresh.

#### Fine-grained state-level alignment (post hoc)

After fuzzy state discovery (Section 3.8) has converged, a state signature is constructed for each state *k* in domain *d*,

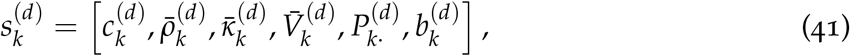

where 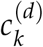 is the state centroid, 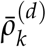 the mean recurrence density, 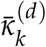 the mean curvature, 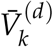 the mean Riemannian volume distortion, 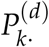 the outgoing state-transition probability vector, and 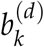 optional biological annotations (e.g. pathway activities, cell-type composition, molecular programmes) when available.

The state-matching cost between state *k* in domain *d* and state *ℓ* in domain *d′* is defined as

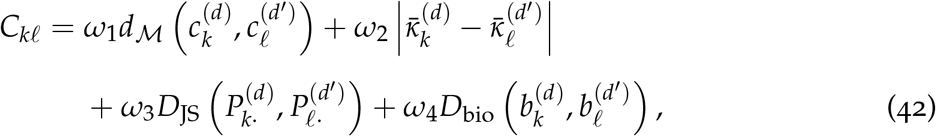

where *D*_JS_(·, ·) denotes the Jensen–Shannon divergence between the state-transition probability distributions and *D*_bio_(·, ·) the cosine distance between biological descriptor vectors. Following [36],

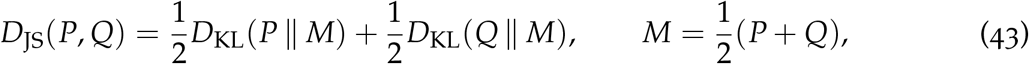

where *D*_KL_(*P* ∥ *M*) = ∑_*ℓ*_ *P*_*ℓ*_ log *P*_*ℓ*_/*M*_*ℓ*_ is the Kullback–Leibler divergence. Unlike *D*_KL_, *D*_JS_ is symmetric and bounded, 0 ≤ *D*_JS_ ≤ log 2, making it a stable, order-independent choice for comparing transition behaviour, and

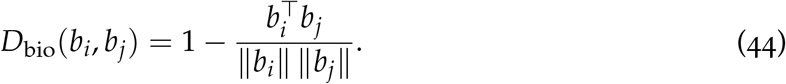

Using this cost, states are aligned across domains by solving a separate entropy-regularised optimal transport (OT) problem [25] over the state signatures,

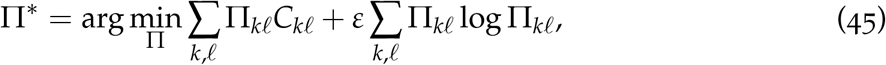

subject to

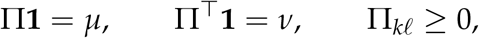

where *µ* and *ν* denote the fuzzy-state mass distributions in domains *d* and *d′*, respectively.

The distinction between coarse alignment and fine-grained state matching is important: the alignment loss ℒ_alignment_ shapes the latent representations during training so domain manifolds become geometrically comparable, whereas the optimal transport plan Π^∗^ is computed only after state discovery converges and provides explicit correspondences between discovered disease states. The two are complementary rather than redundant: coarse alignment establishes a common geometric reference frame, while post hoc state-level alignment identifies biologically corresponding states across domains.

When biological annotations are available for at least a subset of observations, they additionally guide representation learning through a semi-supervised biological consistency loss. Let ℒ ^(*d*)^ ⊆ *{*1, … , *N*_*d*_*}* denote the annotated observations in domain *d*, with annotation 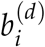 and prediction head *h*_*ω*_,

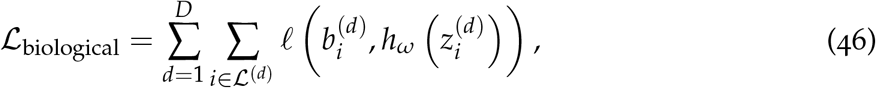

where *ℓ* denotes cross-entropy loss for categorical annotations (e.g. cell type) or squarederror loss for continuous annotations (e.g. pathway activity). For domains without biological annotations, the corresponding summand is omitted.

#### Visualisation of pairwise similarities

For visual comparison across alignment metrics with different numerical ranges, the pairwise GW and state-level OT costs were transformed into similarities using the adaptive exponential kernel

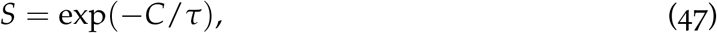

where *C* is the alignment cost and *τ* the median positive off-diagonal cost for each metric. This monotonic transformation preserves the ordering of pairwise relationships while mapping similarities to [0, 1], with larger values indicating greater similarity; it is used solely for visualisation, and all quantitative analyses use the original alignment costs.

### 3.11 Computational Considerations

Two aspects of the algorithm dominate its computational cost: repeated shortest-path computation and repeated optimal-transport solves.

#### Geodesic distances

Exact all-pairs shortest paths over a graph with *N* nodes cost *O*(*N*^2^ log *N*) per refresh using a sparse Dijkstra implementation; recomputing this at every gradient step is prohibitive for tissue-scale data (*N* in the 10^4^–10^5^ range). RGAI mitigates this in three ways: (i) the periodic-refresh schedule of Remark 1, amortising the cost over *T* gradient steps and also reused for curvature (Eq. (25)); (ii) restricting the graph to a sparse *k*-nearest-neighbour or structural-adjacency graph, as already implied by *A*^(*d*)^; and (iii) for very large *N*, a landmark approximation in which exact shortest paths are computed only from a set of *L* ≪ *N* landmark points, with remaining pairwise geodesics estimated via the triangle inequality over landmark distances (landmark multidimensional scaling / landmark Isomap [29]). The same landmark set can be reused as *Ƶ*_land_ in Eq. (37).

#### Optimal transport

Entropic GW solves scale at least quadratically in the number of points aligned. The manifold-level term ℒ_alignment_ therefore operates on a landmark or mini-batch subset (*L* ≪ *N*_*d*_) at each refresh (Section 3.10), while the state-level matching Π^∗^ is solved once, post hoc, over the *K × K* state-signature cost matrix, which is small regardless of *N*.

### 3.12 Joint Optimisation

Although the local metric distance *δ*_*ij*_ is fully differentiable, the global manifold geodesic distance *d*_*M*_, obtained through shortest-path search, is not. RGAI therefore adopts a periodic geometry-refresh strategy: local objective components are evaluated directly from the differentiable edge-local distances *δ*_*ij*_, while global geometric quantities depending on *d*_*M*_ are computed from a precomputed geodesic distance matrix, recomputed every *T* gradient steps and held fixed between refreshes. The same strategy applies to curvature, multiscale recurrence and GW alignment, all of which depend on *d*_*M*_ (Remark 1). This two-timescale procedure enables efficient end-to-end optimisation while avoiding differentiation through the discrete shortest-path operator.

The reconstruction loss is

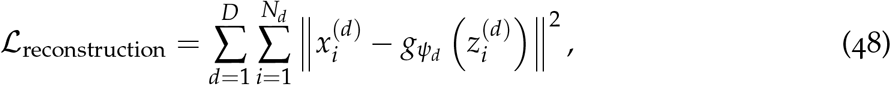

where *g*_*ψd*_ is a domain-specific decoder.

Neighbourhood preservation is encouraged through the fully differentiable, edgelocal objective

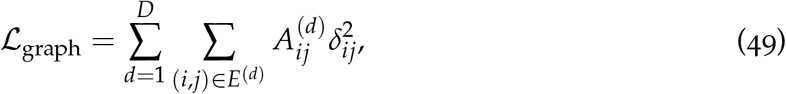

using *δ*_*ij*_ (Eq. (6)) rather than the shortest-path geodesic distance, for the reasons discussed in Remark 1.

With objective components defined (Eqs. (48), (49), (8), (4), (18), (25), (40), and (46)), the overall training objective is

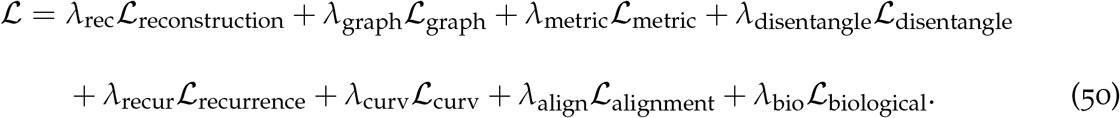

The reconstruction, graph-preservation, metric-learning, disentanglement and biological-supervision components are fully differentiable and provide the gradient-based optimisation used to learn the latent representations and Riemannian metric throughout training. The recurrence-derived quantities ℒ_recurrence_, ℒ_curv_, and ℒ_alignment_ are updated only after each periodic geometry refresh, when the multiscale recurrence tensors and associated descriptors are recomputed from the current latent manifold, and remain fixed between refreshes—avoiding repeated construction of the expensive geodesic recurrence representation while providing periodically updated geometric evaluation.

After optimisation, the framework produces a shared disease-state manifold, fuzzy tissue-state memberships, multiscale recurrence tensors, local curvature and metric maps, state-transition networks, and cross-domain tissue-state correspondences Π^∗^.

The complete procedure is summarised in Algorithms 1 and 2, reflecting the two-timescale strategy above. Algorithm 1 performs the differentiable representation-learning stage, jointly training the domain-specific encoders, shared encoder, metric network and, where available, the biological-annotation head, interleaving gradient-based optimisation with periodic geometry refreshes that recompute geodesic distances, recurrence tensors, curvature descriptors and manifold-level alignment quantities. Algorithm 2 then runs once, after the latent representation converges, performing the fuzzy tissue-state discovery of Section 3.8, the transition analysis of Section 3.9, and the state-level optimal-transport alignment of Section 3.10, producing the final cross-domain tissue-state correspondences.

#### Algorithm 1 RGAI: Representation, Metric, and Recurrence Learning

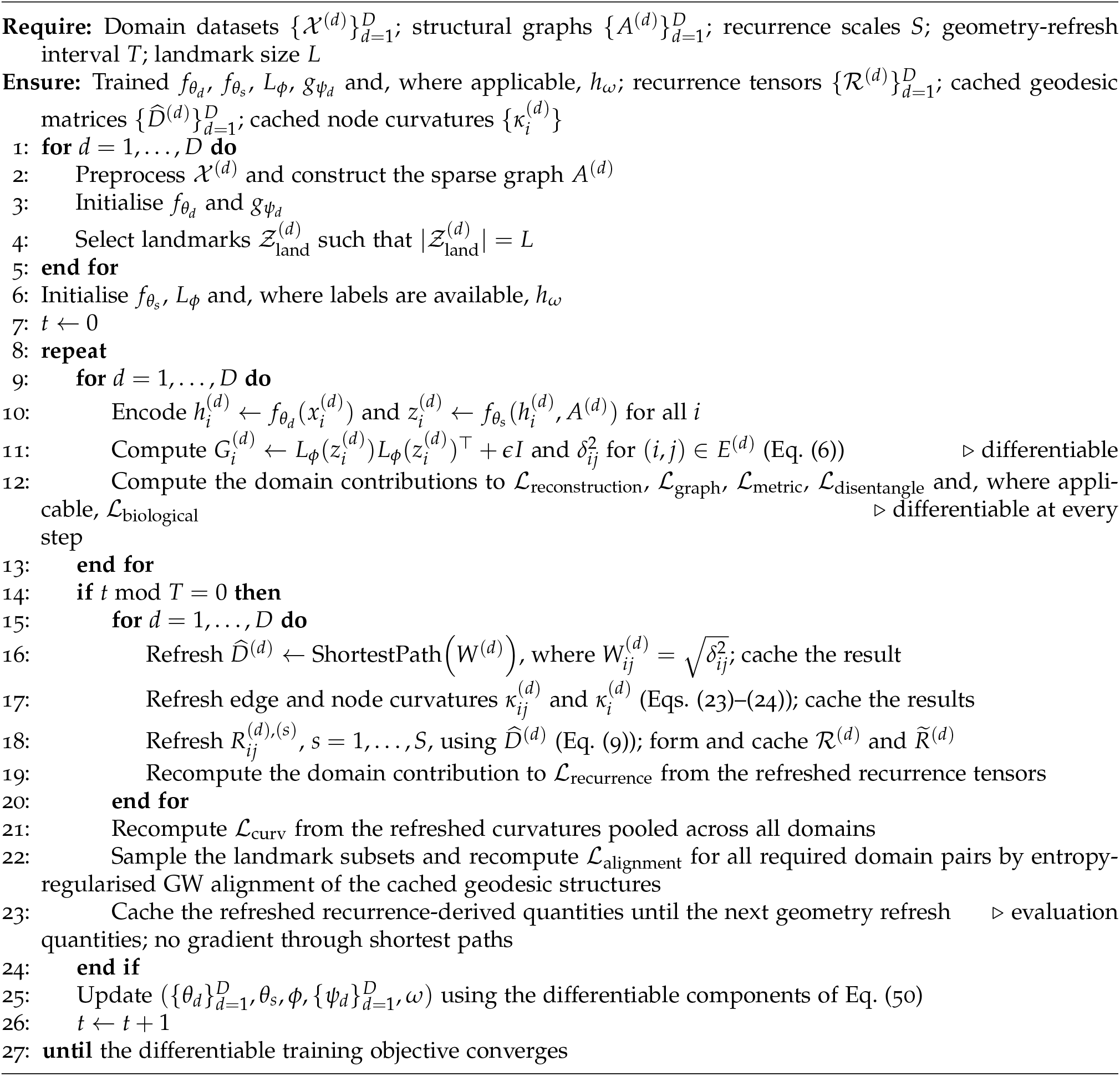

#### Algorithm 2 RGAI: Fuzzy State Discovery and Cross-Domain Alignment

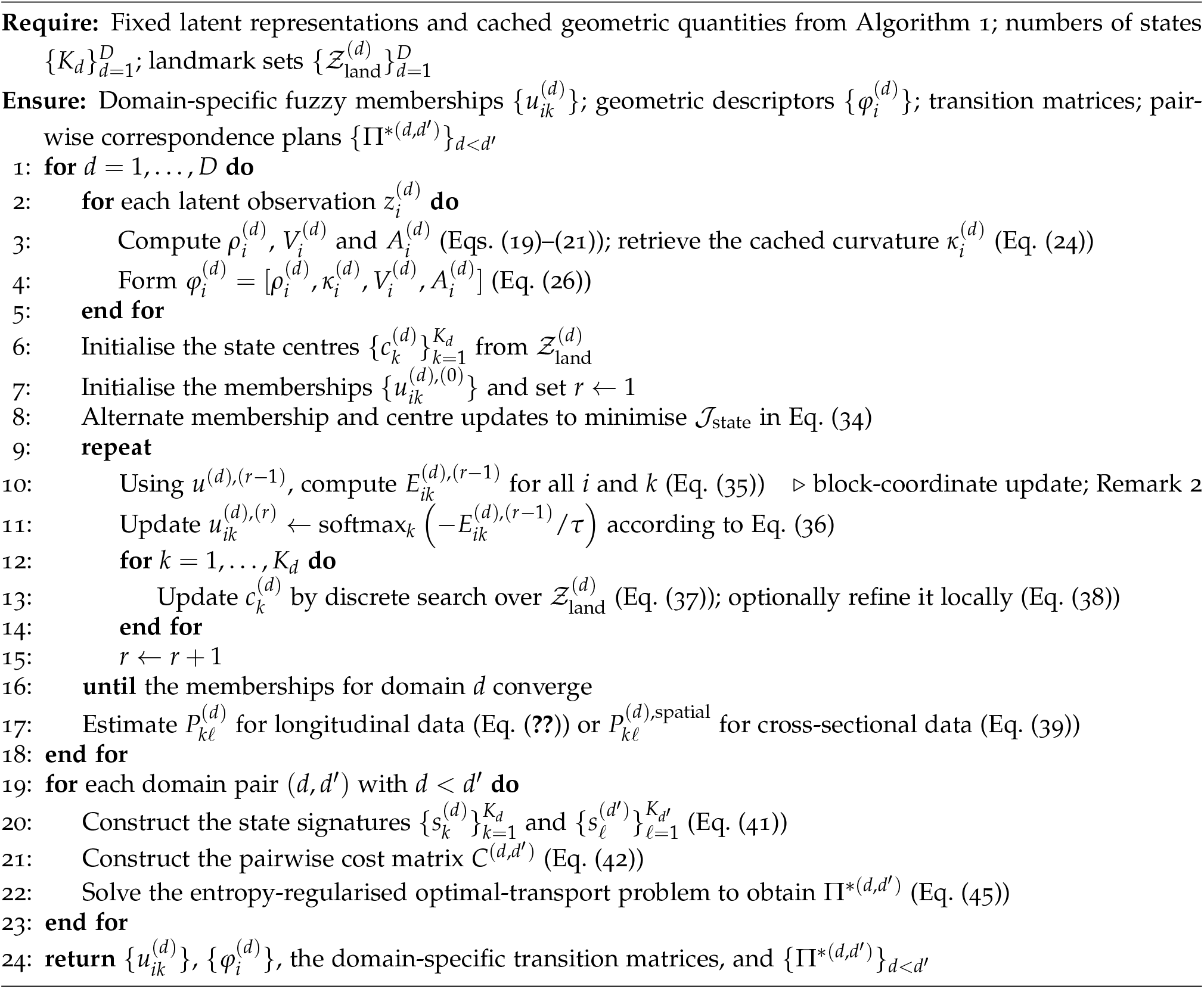

## 4 Results

RGAI was evaluated using the six publicly available 10x Visium spatial transcriptomic datasets described in Section 3.1, spanning tissue repair, chronic inflammatory disease, cardiovascular tissue, and epithelial cancers. Unless otherwise stated, experiments used the common hyperparameter settings in Table 2, with identical configurations across datasets for direct cross-domain comparison. Results are organised by the key components of the framework rather than by dataset.

**Table 2.** Common hyperparameter settings used for all datasets.

| Parameter | Value | Rationale |
| --- | --- | --- |
| Hidden dimension ($d_{\text{hidden}}$) | 64 | Provides sufficient representational capacity while avoiding excessive model complexity. |
| Shared latent dimension | 10 | Captures common biological structure shared across domains. |
| Domain-specific latent dimension | 6 | Preserves domain-specific variation not shared across tissues. |
| Learning rate | $10^{-3}$ | Standard stable setting for Adam optimisation. |
| Maximum training iterations | 500 | Sufficient for convergence across all analysed datasets. |
| Geometry refresh interval ($T$) | 10 | Balances adaptation of the learned geometry against computational cost. |
| Spatial nearest neighbours ($k$) | 6 | Preserves local tissue topology while maintaining graph connectivity. |
| Shared variable genes | 2000 | Widely adopted compromise between retaining informative biological variation and reducing noise and computational cost. |
| Recurrence scales ($\alpha_s$) | $\{0.5, 1, 2\}$ | Capture fine-, intermediate-, and coarse-scale recurrence structure. |
| Recurrence exponent ($q$) | 2 | Produces a smooth Gaussian-like decay of recurrence with distance. |
| Metric regularisation ($\epsilon$) | $10^{-3}$ | Ensures numerical stability and positive definiteness of the learned metric tensor. |
| Landmarks ($L$) | 75 | Reduces the computational cost of GW alignment while preserving global manifold structure. |
| Candidate numbers of states ($K$) | 3–8 | Covers the expected range of tissue-state complexity for automatic model selection. |
| Fuzzy exponent ($m$) | 2 | Standard setting in fuzzy c-means, balancing crisp and diffuse memberships. |
| OT regularisation ($\zeta$) | 0.05 | Provides stable entropy-regularised transport plans without excessive smoothing. |
| GW regularisation ($\eta$) | 0.05 | Stabilises GW optimisation while preserving meaningful geometric correspondences. |
| Random seed | 42 | Ensures reproducibility of all experiments. |
| Independent runs | 10 | Assesses robustness to random initialisation. |

### 4.1 Automatic determination of latent tissue-state complexity

The multi-criterion model-selection procedure consistently identified well-supported optimal latent tissue-state complexities across the six datasets (Table 3). Five datasets (wound healing, periodontitis, HNSCC, cardiac tissue and CRC) were optimally represented by three latent tissue states, whereas OSCC required four, indicating more complex organisation than the other domains.

**Table 3.** Automatic determination of latent tissue-state complexity: selected *K*, aggregate model-selection score, second-best candidate, score margin, and qualitative organisation for each dataset. Lower scores indicate better-supported solutions. GH = greater heterogeneity.

| Dataset | Selected $K$ | Selection score | Second-best $K$ | Score margin | Organisation |
| --- | --- | --- | --- | --- | --- |
| Wound healing | 3 | 12 | 4 | 1 | Compact |
| Periodontitis | 3 | 5 | 4 | 9 | Compact |
| OSCC | 4 | 12 | 3 | 1 | GH |
| HNSCC | 3 | 7 | 4 | 7 | Compact |
| Cardiac | 3 | 12 | 5 | 3 | Compact |
| CRC | 3 | 10 | 4 | 3 | Compact |

The aggregate selection scores and margins supported these choices: periodontitis and HNSCC showed the largest margins over the second-best candidate, indicating strong separation, while wound healing and OSCC showed smaller but still positive margins. The selection-score profiles in Fig. 1 exhibited a distinct optimum for every dataset, indicating that the selected solutions were supported by multiple complementary criteria rather than a monotonic preference for increasing complexity.

**Figure 1.**
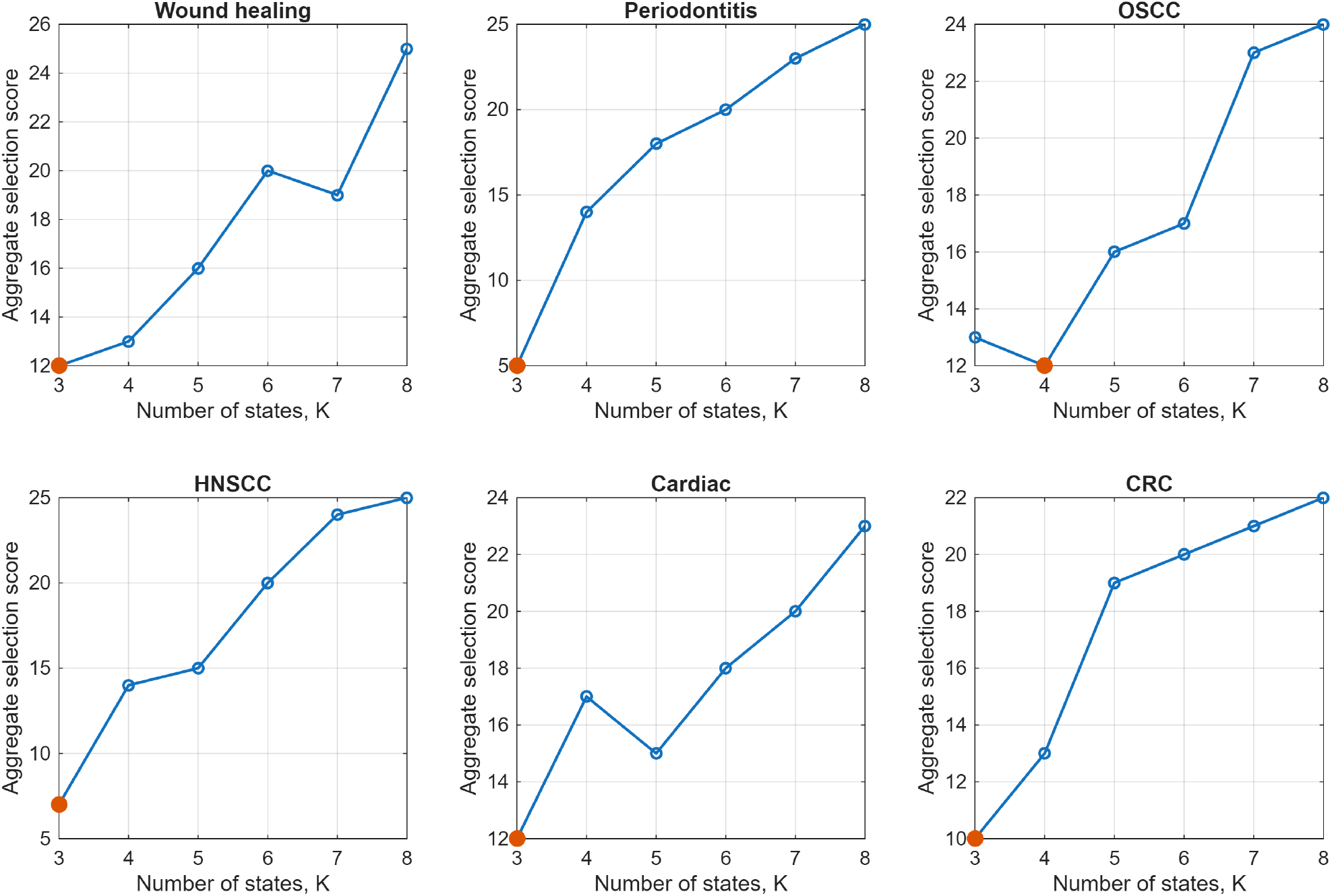
Automatic determination of latent tissue-state complexity. Aggregate model-selection scores for candidate numbers of latent tissue states; filled markers indicate the selected solution. Five datasets selected three states, whereas OSCC selected four, indicating structural heterogeneity.

### 4.2 Learned recurrence geometry of tissue states

The learned latent manifolds exhibited distinct geometric characteristics across the six domains, indicating that the learned Riemannian geometry adapted to the latent organisation of each tissue. Figure 2 summarises the distributions of four complementary descriptors: local recurrence density, latent manifold curvature, local geometric anisotropy, and logvolume distortion.

**Figure 2.**
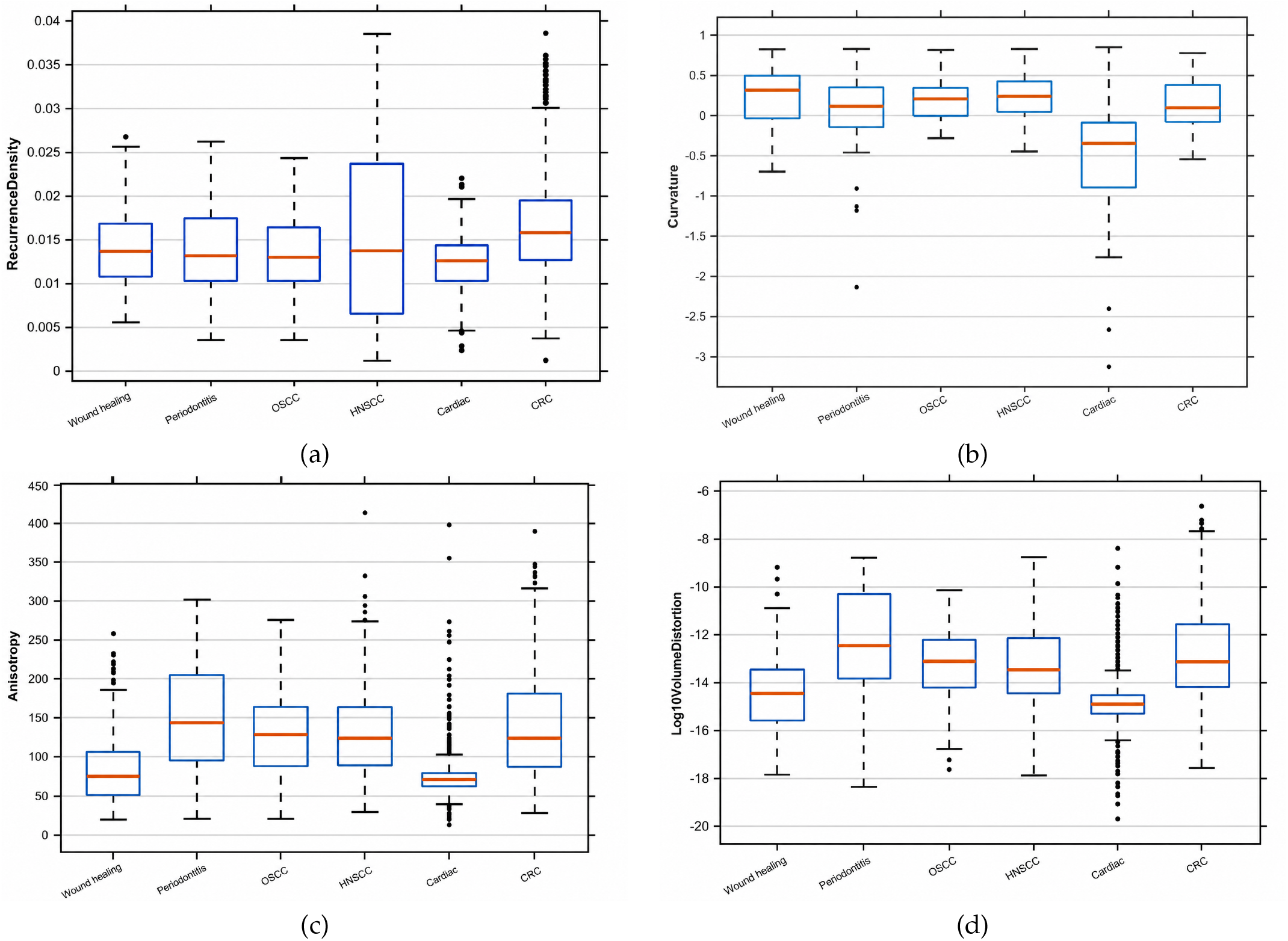
Learned recurrence geometry across the six domains: (a) local recurrence density, (b) latent manifold curvature, (c) local geometric anisotropy, (d) log-volume distortion. Differences among domains indicate distinct recurrence-geometric organisations and provide signatures for cross-domain alignment.

Local recurrence density (Fig. 2(a)) varied substantially among datasets, with wound healing, periodontitis, HNSCC and CRC showing greater variability than cardiac tissue. Curvature (b) also differed markedly, with both positive and negative values observed and the spread differing among datasets. Anisotropy (c) likewise differed across domains, from compact to broader distributions extending toward larger values, and log-volume distortion (d) differed among datasets, showing that the learned metric expanded or contracted local regions to varying degrees across tissues.

### 4.3 Relationships among learned geometric descriptors

To investigate relationships among the learned geometric quantities, pairwise Spearman correlations were computed among local recurrence density, curvature, anisotropy, and tissue-state membership confidence within each domain (Fig. 3). Associations varied substantially across the six datasets rather than following a common structure, indicating domain-specific interactions among the descriptors. The strongest associations were generally between curvature and anisotropy and between curvature and recurrence density, though magnitude and direction differed among tissues.

**Figure 3.**
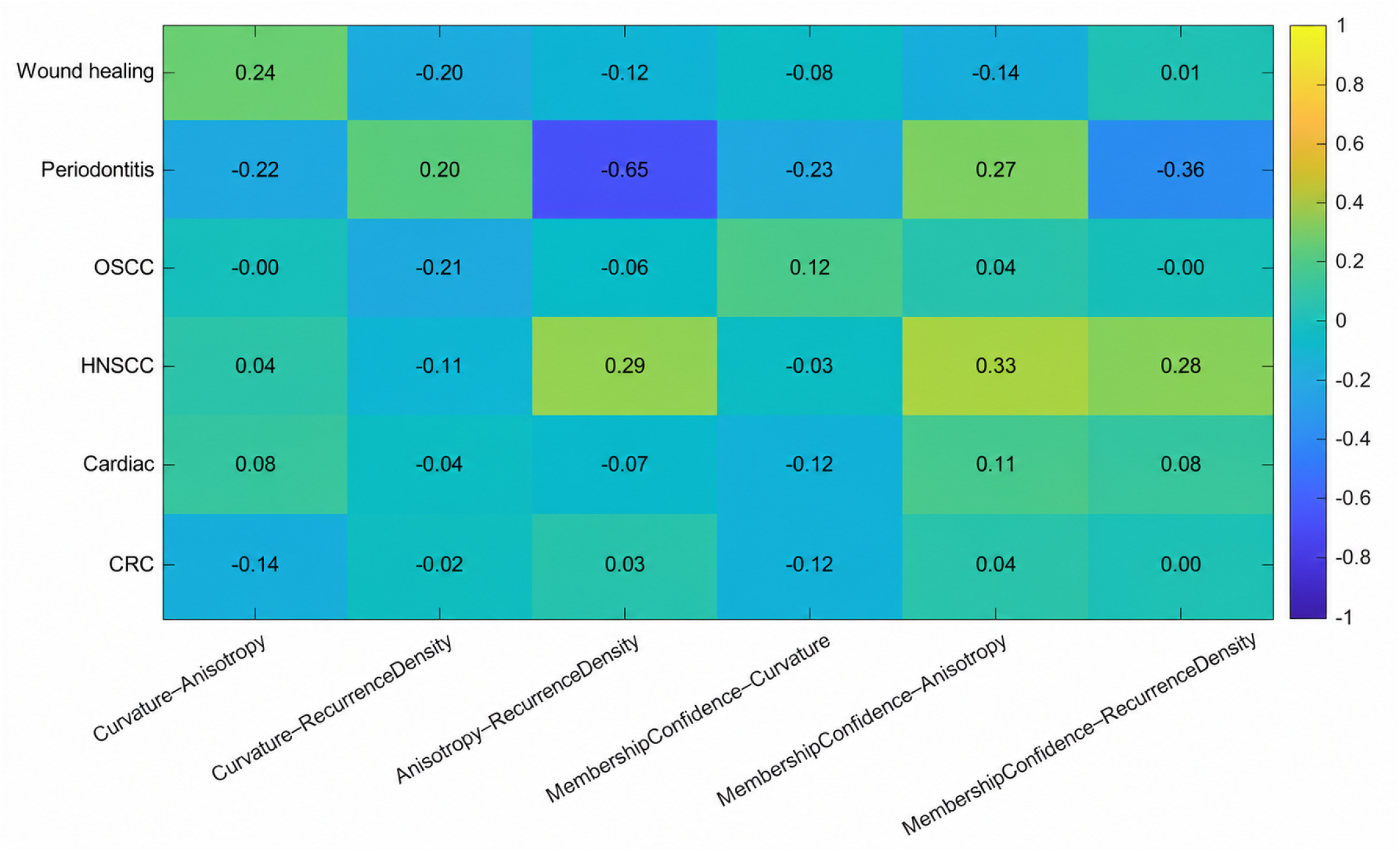
Within-domain associations among learned tissue-state descriptors. Each row is one dataset; each column reports the Pearson correlation between two descriptors computed within that dataset (curvature, anisotropy, recurrence density, fuzzy membership confidence), demonstrating complementary rather than redundant characteristics prior to cross-domain alignment.

Overall, the heterogeneous correlation patterns suggest that the descriptors capture complementary rather than redundant aspects of latent tissue-state organisation, and that no single descriptor adequately characterises it; each dataset instead exhibited its own characteristic combination.

### 4.4 Cross-domain geometric alignment and tissue-state correspondence

Pairwise GW alignment was performed to quantify the similarity of learned latent manifold geometries across the six domains. The optimisation converged for most domain pairs, and the remainder reached the maximum permitted iterations with negligible further change, indicating stable numerical behaviour (Table 4).

**Table 4.** Pairwise GW optimisation statistics: total inner transport iterations, outer optimisation iterations, and final outer residual for each domain pair.

| Domain pair | Inner iterations | Outer iterations | Outer residual |
| --- | --- | --- | --- |
| Wound healing–Periodontitis | 105 | 47 | $8.97 \times 10^{-5}$ |
| Wound healing–OSCC | 45 | 16 | $7.56 \times 10^{-5}$ |
| Wound healing–HNSCC | 240 | 18 | $6.85 \times 10^{-5}$ |
| Wound healing–Cardiac | 125 | 14 | $6.98 \times 10^{-5}$ |
| Wound healing–CRC | 55 | 13 | $4.90 \times 10^{-5}$ |
| Periodontitis–OSCC | 100 | 12 | $7.67 \times 10^{-5}$ |
| Periodontitis–HNSCC | 115 | 9 | $5.63 \times 10^{-5}$ |
| Periodontitis–Cardiac<sup>†</sup> | 1000 | 17 | $8.98 \times 10^{-5}$ |
| Periodontitis–CRC | 125 | 24 | $9.68 \times 10^{-5}$ |
| OSCC–HNSCC | 75 | 9 | $4.32 \times 10^{-5}$ |
| OSCC–Cardiac | 625 | 20 | $9.16 \times 10^{-5}$ |
| OSCC–CRC | 40 | 23 | $8.18 \times 10^{-5}$ |
| HNSCC–Cardiac | 460 | 29 | $9.91 \times 10^{-5}$ |
| HNSCC–CRC | 90 | 15 | $8.65 \times 10^{-5}$ |
| Cardiac–CRC | 195 | 29 | $8.56 \times 10^{-5}$ |

<sup>†</sup> Reached the maximum permitted inner iterations before satisfying the stopping criterion; the final residual ( $8.98 \times 10^{-5}$ ) was comparable to other alignments, indicating a stable solution.

The pairwise GW similarities are summarised in Fig. 5(a). Considerable variability was observed: several domain pairs showed relatively high similarity despite representing distinct biological systems, while others showed substantially weaker similarity, indicating that RGAI captures both shared and domain-specific patterns of latent organisation without prior biological correspondences. Comparing GW with state-level OT similarities (Fig. 4) showed broadly similar but non-identical trends: domain pairs with similar global geometry did not always show equally strong tissue-state correspondence, indicating the two measures capture complementary aspects of cross-domain organisation, as further illustrated by the similarity matrices in Fig. 5.

**Figure 4.**
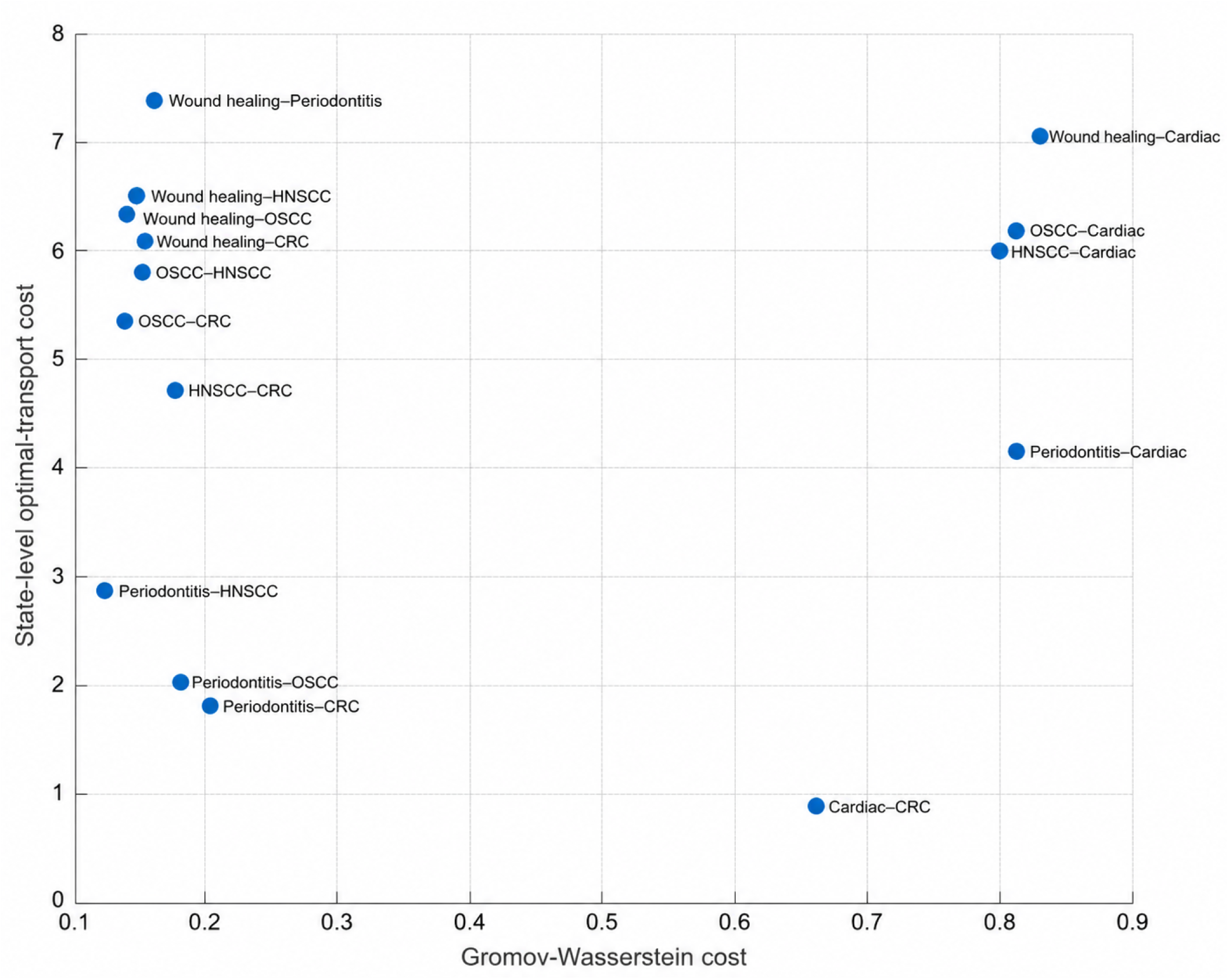
Coarseand fine-scale cross-domain alignment. Each point is one domain pair; the horizontal axis shows GW alignment cost (coarse-scale manifold similarity) and the vertical axis shows state-level OT cost (fine-scale tissue-state correspondence). The dispersion shows that similar global geometry does not necessarily imply similar tissue-state correspondence.

**Figure 5.**
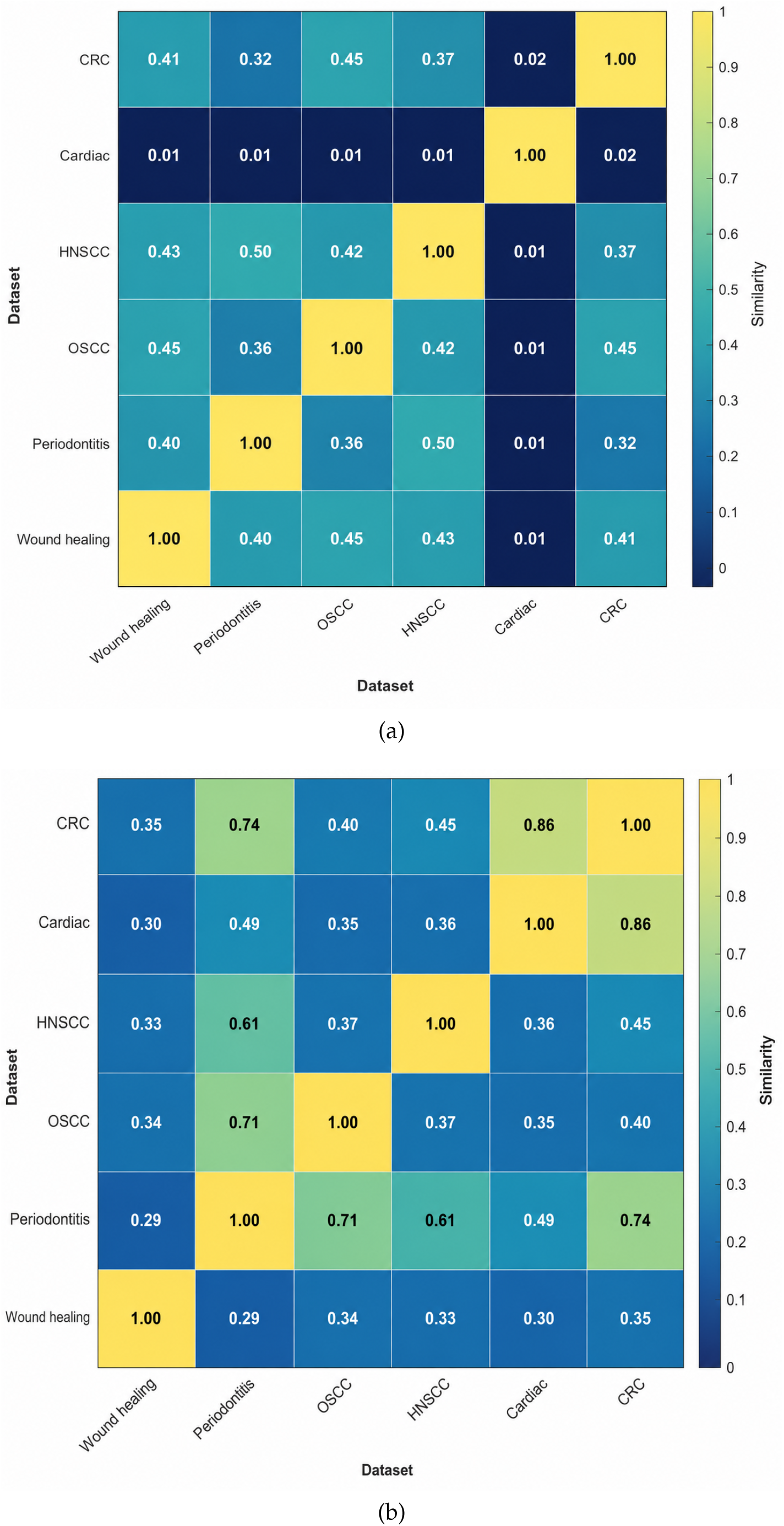
Cross-domain geometric similarity and tissue-state correspondence. (a) Pairwise GW similarities and (b) pairwise state-level OT similarities, both obtained via the transformation in Eq. (47). Unlike Fig. 4, which shows raw costs, these heatmaps show transformed similarities for direct visual comparison.

## 5 Discussion

### 5.1 RGAI learns biologically meaningful recurrence geometry

A principal contribution of RGAI is the explicit learning of recurrence geometry through a data-adaptive Riemannian metric, rather than a fixed metric imposed on latent similarity— particularly relevant for spatial transcriptomic data, where tissue organisation often exhibits gradual transitions, branching structures and heterogeneous spatial densities unlikely to be captured by a single global metric. A related advantage is the automatic determination of latent tissue-state complexity: instead of specifying the number of states a priori, the multi-criterion model-selection procedure chose the complexity best supported by the data, with five datasets optimally represented by three states and OSCC by four, reducing dependence on user-defined parameters while letting the learned representation reflect each domain’s characteristics.

The learned recurrence geometries support this adaptivity: across the six datasets, the distributions of recurrence density, curvature, anisotropy and log-volume distortion differed substantially, and the heterogeneous correlation patterns among these descriptors suggest that they capture complementary rather than redundant aspects of latent tissue-state organisation—more appropriately characterised by multiple interacting geometric descriptors than by any single quantity. Because geodesic distances are computed with respect to the learned local geometry of each latent manifold, the subsequent GW alignment compares domains according to their learned manifold geometry rather than absolute latent coordinates, enabling comparison even when different tissues occupy distinct regions of the latent space. Overall, these findings indicate that integrating learned Riemannian geometry, recurrence analysis and optimal transport provides a coherent framework for analysing heterogeneous spatial transcriptomic data.

### 5.2 Conserved recurrence organisation across distinct diseases

Although the analysed datasets represent different organs, disease processes and experimental studies, cross-domain alignment identified measurable similarities in their learned latent manifold geometry, with some domain pairs showing relatively strong GW correspondence and others substantially weaker similarity—indicating that the learned recurrence geometries contain both shared and domain-specific patterns of tissue-state organisation. This is noteworthy because the alignment is based on the organisation of the learned manifolds rather than direct comparison of gene-expression profiles, so identified correspondences reflect geometric similarity even where the underlying biology differs substantially; the complementary state-level OT analysis further showed that similarity of global geometry does not necessarily imply identical tissue-state correspondence.

These findings suggest that distinct tissues may share aspects of their latent geometric organisation despite differences in cellular composition and molecular context, possibly reflecting common patterns of tissue remodelling, inflammation or repair, though biological validation is required. Domain pairs with similar global geometry but dissimilar state correspondence may share broad organisational principles while differing in molecular programmes, whereas domains with distinct geometries but similar state correspondence may contain conserved pathological states despite differing architecture—a hierarchical perspective that could inform mechanism-based disease stratification and biomarker discovery, though these remain hypotheses requiring validation in independent cohorts.

### 5.3 Complementary roles of manifold alignment and state-level optimal transport

The separation of cross-domain comparison into two complementary stages—GW alignment of the learned manifolds, followed by entropy-regularised OT between tissue-state signatures—lets similarity be examined at both the global and finer tissue-state levels. As shown in Section 4, the two measures followed broadly similar but non-identical trends, confirming that manifold-level and tissue-state organisation capture related but complementary characteristics of biological structure, together offering a more comprehensive basis for cross-domain analysis.

### 5.4 Advantages over existing cross-domain spatial transcriptomic integration methods

Existing methods for integrating spatial transcriptomic datasets are primarily designed to reduce batch effects, align shared cell populations, or construct common latent representations, and have proved valuable when substantial biological correspondence exists among the analysed datasets; comparative analysis across different organs and disease processes is more challenging, however, because the organisation of latent tissue states may differ substantially between domains. RGAI addresses this differently: rather than constructing a single shared latent representation, it first learns a domain-specific latent manifold with its associated Riemannian geometry, then performs cross-domain comparison via GW alignment, letting each domain retain its own geometric organisation while enabling comparison through relationships among the learned manifolds rather than direct coordinate correspondence.

RGAI also produces a set of interpretable geometric descriptors—recurrence density, curvature, anisotropy and log-volume distortion—that characterise complementary properties of the learned manifolds and, together with the alignment analyses, provide a geometric perspective on tissue organisation complementing conventional analyses based on molecular similarity or latent embeddings. The present study demonstrates the feasibility of this framework across six datasets spanning multiple organs and disease processes; systematic benchmarking against established integration methods on larger, more diverse dataset collections is an important direction for future work.

#### Potential clinical and translational implications

Cross-domain analysis may help identify shared pathological mechanisms not apparent when diseases are studied independently. By distinguishing conserved tissue-state organisation from domain-specific alterations, RGAI could support more reproducible biomarkers, mechanism-based patient stratification, and prioritisation of therapeutic targets relevant across diseases, and shared tissue-state programmes may motivate drug repurposing or cross-disease basket trials. These applications require validation in clinically annotated, prospective cohorts; the present analysis is a framework for generating translational hypotheses rather than a clinical decision-support tool.

### 5.5 Limitations

Several limitations should be acknowledged. Although RGAI was evaluated across six datasets spanning multiple tissues and disease processes, a larger and more diverse collection will be required to establish generality, including robustness across additional platforms, species, tissue types and protocols. The current implementation performs pairwise GW alignment between domains; while this enables detailed, interpretable pairwise comparison, extending the framework to jointly align multiple domains may improve scalability for large atlases. The biological interpretation of the learned descriptors also remains largely inferential: the underlying mechanisms remain to be established experimentally, requiring complementary molecular measurements (spatial proteomics, multiplex imaging, longitudinal observations) together with targeted validation. Finally, this study focused on demonstrating feasibility rather than benchmarking against existing integration methods; systematic comparative evaluation across a broad range of datasets and tasks is needed to establish the framework’s practical strengths and limitations.

### 5.6 Future directions

RGAI opens several opportunities for future research. One natural extension is joint multidomain manifold alignment, enabling multiple tissues and disease conditions to be analysed simultaneously rather than through pairwise comparisons, facilitating large-scale atlases organised by learned geometric relationships. Another is integrating complementary molecular and imaging modalities—spatial proteomics, multiplex imaging, metabolomics and longitudinal spatial transcriptomics—which may give a more comprehensive representation of tissue-state organisation.

The study also motivates further theoretical development: RGAI extends concepts from fuzzy recurrence analysis and recurrence networks [37, 38] to spatially organised transcriptomic data, and future work could pursue more general formulations representing spatial organisation, temporal dynamics and multimodal data within a common framework. Methodologically, uncertainty quantification, adaptive metric learning, scalable optimal-transport algorithms and more efficient optimisation may improve robustness and scalability for larger datasets, alongside more extensive benchmarking and experimental validation. Finally, the framework may be applicable beyond spatial transcriptomics—to single-cell sequencing, histopathology, medical imaging and longitudinal disease studies— offering a common geometric framework for heterogeneous biological systems.

## 6 Conclusion

This study introduced RGAI, a framework for cross-domain tissue-state analysis integrating learned Riemannian geometry, recurrence analysis, GW manifold alignment and entropy-regularised OT. By learning domain-specific latent geometry and deriving recurrence-based geometric descriptors—recurrence density, curvature, anisotropy and log-volume distortion—RGAI provides an interpretable representation of tissue-state organisation across heterogeneous spatial transcriptomic datasets, with a hierarchical alignment strategy combining manifold-level geometric comparison with state-level tissue correspondence.

The results show that RGAI automatically adapts latent tissue-state complexity to individual datasets, learns distinct recurrence geometries across biological domains, and identifies both shared and domain-specific patterns of latent manifold organisation through cross-domain alignment. More broadly, RGAI extends previous work on fuzzy recurrence analysis and recurrence networks to learned geometric representations, bringing together recurrence analysis, differential geometry and optimal transport within a single AI framework. Although developed and evaluated for spatial transcriptomics, the underlying formulation may also prove useful for other high-dimensional biomedical data in which relationships among observations are more naturally characterised by learned geometric organisation than by direct feature similarity.

## Code Availability

The MATLAB package implementing the proposed framework is publicly available at the author’s website: https://sites.google.com/view/tuan-d-pham/codes, under the heading “Recurrence Geometric AI in Spatial Transcriptomics”.

## Notes

https://www.ncbi.nlm.nih.gov/geo/query/acc.cgi?acc=GSE241124

https://www.ncbi.nlm.nih.gov/geo/query/acc.cgi?acc=GSE208253

https://www.ncbi.nlm.nih.gov/geo/query/acc.cgi?acc=GSE181300

https://www.ncbi.nlm.nih.gov/geo/query/acc.cgi?acc=GSE135805

https://www.ncbi.nlm.nih.gov/geo/query/acc.cgi?acc=GSE226997

### Competing Interest Statement

The authors have declared no competing interest.

